# Mechanistic insights into the activity of SARS-CoV-2 RNA polymerase inhibitors using single-molecule FRET

**DOI:** 10.1101/2024.10.15.618435

**Authors:** Danielle Groves, Rory Cunnison, Andrew McMahon, Haitian Fan, Jane Sharps, Adrian Deng, Jeremy R. Keown, Ervin Fodor, Nicole C. Robb

**Affiliations:** Warwick Medical School, University of Warwick, Coventry, CV4 7AL, UK; Sir William Dunn School of Pathology, University of Oxford, Oxford, OX1 3PU, UK; School of Life Sciences, University of Warwick, Coventry, CV4 7AL, UK

## Abstract

The COVID-19 pandemic, caused by the SARS-CoV-2 virus, has resulted in significant global mortality and disruption. Despite extensive research, the precise molecular mechanisms underlying SARS-CoV-2 replication remain unclear. To address this, we developed a single-molecule Förster resonance energy transfer (smFRET) assay to directly visualize and analyse in vitro RNA synthesis by the SARS-CoV-2 RNA-dependent RNA polymerase (RdRp). We purified the minimal replication complex, comprising nsp12, nsp7, and nsp8, and combined it with fluorescently labelled RNA substrates, enabling real-time monitoring of RNA primer elongation at the single-molecule level. This platform allowed us to investigate the mechanisms of action of key inhibitors of SARS-CoV-2 replication. In particular, our data provides evidence for remdesivir’s mechanism of action, which involves polymerase stalling and subsequent chain termination dependent on the concentration of competing nucleotide triphosphates. Our study demonstrates the power of smFRET to provide dynamic insights into SARS-CoV-2 replication, offering a valuable tool for antiviral screening and mechanistic studies of viral RdRp activity.

## Introduction

The emergence of the novel SARS-CoV-2 virus at the end of 2019 has resulted in the ongoing COVID-19 pandemic that has infected millions of people and caused worldwide social and economic disruption. In spite of the devastating impact of COVID-19, many questions regarding the exact molecular details of how coronaviruses replicate remain unanswered and hence the development of intervention strategies to combat SARS-CoV-2 infection relies on increasing our understanding of the basic mechanisms of viral replication. To date, much of our knowledge of SARS-CoV-2 replication has relied on data obtained using ensemble methods that report on the mean properties of billions of molecules, averaging the measured parameters over the entire molecular population. Single-molecule techniques allow real-time studies of viral replication with the advantage that they can provide direct observations of just one molecule at a time.

The SARS-CoV-2 virus contains a 30kb single-stranded positive-sense genome that is replicated by the viral RNA-dependent RNA polymerase (RdRp). Several cryo-EM studies of the SARS-CoV-2 RdRp alone or bound to RNA have elucidated the basic structure of the complex^1–3^. The complex consists of three types of non-structural proteins (nsps): a single copy of nsp12, which contains the RdRp active site and the nidovirus RdRp-associated nucleotidyltransferase (NiRAN) domain which acts as a capping enzyme^4^, a single copy of nsp7, suggested to be responsible for stabilisation of regions of nsp12 thought to be involved in RNA binding^5^, and two copies of nsp8, proposed to be essential for processivity of the polymerase^1,2^. It is believed that nsp12, nsp7 and nsp8 represent a minimal replication transcription complex (the RTC) capable of replicating RNA, making the complex an attractive target for antiviral drug development^3^.

Comparison of the apo form of SARS-CoV-2 RdRp^2^ with structures bound to RNA^5,6^ showed a conserved structure for the RdRp which does not change upon RNA binding. Cryo-EM and biochemical studies of the replicating RdRp^3,5^ show that during replication nuclear triphosphates (NTPs) enter the entry channel and are then stabilised in the active site of nsp12 by hydrogen-bonding, during which the RdRp adopts the ‘pre-translocation’ state. The complex adopts a ‘post-translocation’ state when primer RNA enters the +1 position within the RNA-binding cleft. The polymerase domain of nsp12 contains aspartate residues D760 and D761, which bind to two Mg^2+^ ions during translocation, and are thus essential for the catalysis of nucleotide addition^3,5^. Watson-Crick base pairing between template and product RNA is completed by a condensation reaction which stabilises the post-translocation conformation of RdRp. The nsp8 pair form positive charges either side of the active site with long helical ‘sliding poles’ which pull negatively charged RNA through the complex^2,3^.

Structural insights into the SARS-CoV-2 RdRp have allowed the function of a variety of non-nucleoside or nucleoside analogue inhibitors to be explored. For example, remdesivir is a broad-spectrum antiviral medicine that was originally developed to treat hepatitis C^7^ and filoviruses such as Ebola^8^ before being used as a post-infection treatment for COVID-19. Remdesivir is an adenosine analogue that competes with the natural adenosine triphosphate (ATP) substrate to inhibit viral RdRps. The primary mechanism of inhibition has been suggested to be the incorporation of remdesivir triphosphate into nascent viral RNA by the viral RdRp, resulting in chain termination^9–11^. Unlike many other chain terminators, this is not mediated by preventing addition of the immediately subsequent nucleotide, but is instead delayed, occurring after additional bases have been added to the growing RNA chain. For MERS-CoV, SARS-CoV-1, and SARS-CoV-2, arrest of RNA synthesis occurs after incorporation of three additional nucleotides^10,11^. Because of read-though following remdesivir incorporation, the analogue may be incorporated into the viral RNA template strand, which has also been shown to also block RNA synthesis via a second, template-dependent inhibition mechanism^12^. Suramin is an example of a non-nucleoside inhibitor that has been shown to bind directly to the SARS-CoV-2 RdRp, and results in inhibition by blocking the binding of the RNA template strand and preventing entry of product RNA into the active site^13^.

Here, we have used a solution-based single-molecule Förster Resonance Energy Transfer (smFRET) assay to measure the dynamic conformational changes that occur during SARS-CoV-2 RdRp-mediated RNA extension and inhibition. We have shown that we can reconstitute minimal viral replication complexes in vitro and monitor the conformational changes that take place in RNA during RdRp-mediated extension via the movement of fluorescence dyes positioned on the RNA. We have applied this approach to probe the mechanisms of action of both a nucleoside and non-nucleoside inhibitor of SARS-CoV-2, validating their inhibition and differentiating between different mechanisms of action. This technique is therefore well suited to deciphering the mechanisms involved in viral replication inhibition, and we anticipate can be further applied to the screening of multiple other viral polymerases and inhibitors.

## Methods

### RNA

We ordered minimal RNA oligonucleotides from Integrated DNA Technologies (IDT) based on the sequences in a cryo-EM structure of SARS-CoV-2 RdRp^14^, conjugated to fluorescent labels Cy3, Cy5 and ATTO647 via 6-carbon linkers. The sequences and positions of the fluorescent dyes are described in the figures and in Supplementary Table 1. Template and primer RNAs were annealed at a final concentration of 300 nM in hybridization buffer (50 mM Tris-HCl pH 8.0, 1 mM EDTA, 500 mM NaCl) using a single cycle temperature gradient from 95 - 4°C.

### Protein

The SARS-CoV-2 RdRp wild type (WT) nsp12 and active site mutant nsp12 D760A/D761A were cloned into a MultiBac plasmid with a Tobacco Etch Virus (TEV) protease cleavable protein-A tag^15^ and a histidine tag at the C-terminus, whilst the plasmids for nsp7 and nsp8 were transformed into BL21 E. coli using a lac promoter and then expressed. The expression and purification of nsps has been published previously^4,16^. Briefly, for nsp12 and nsp12 D760A/D761A, 0.5 million cells/mL Sf9 insect cell culture (Sf-900II medium, Life Technologies) was infected with 5 mL V1 baculovirus (MOI = 1). The cell culture was harvested 3 days post infection, then flash frozen with liquid nitrogen and stored at -80°C. For nsp7 and nsp8, E. coli was cultured overnight from glycerol stocks in LB media, then inoculated into a greater volume of LB for overnight cell culture. Cells were induced with 0.5 mM Isopropyl β-D-1-thiogalactopyranoside (IPTG) when OD600 = 0.6-0.8, then the cell culture was grown at 18°C overnight before harvesting by centrifugation at 3500g/15 minutes.

1-2 L of sf9 or E. coli pellet(s) was resuspended in wash buffer (50 mM HEPES-NaOH pH7.5, 300 mM NaCl, 10% Glycerol, 0.05% OTG, 1mM DTT) with protease inhibitor before cell lysis by sonication. The cell lysate was spun down and the supernatant was incubated with 1mL IgG or GST Sepharose per litre of cell culture for 3-4 hours at 4°C. The solution was washed 3 times with wash buffer, then the protein A tag was cleaved with TEV protease (0.5-1 mg TEV / L cell culture, in the presence of 1 mM DTT), or the GST tag was cleaved with 3C protease (0.2-0.5 mg 3C / L cell culture, in the presence of 1 mM DTT), overnight at 4°C. The next day, the supernatant (tag-cleaved protein) was concentrated to 0.5-2 mL and loaded on to an AKTA Pure Superdex 200 with 10/300 increase column for nsp12 and nsp12 D760A/D761A and Superdex 75 for nsp7 and nsp8 in SEC running buffer (25 mM HEPES-Na (pH 7.5), 100 mM NaCl, 2 mM MgCl2, 1 mM DTT). The fractions containing the target protein were pooled and checked by SDS-PAGE against the pre-stained precision plus BioRad ladder 10–250 kDa (161-0373), then concentrated to ∼ 5-10 mg/mL with Amicon Millipore MWCO concentrators using a 100 kDa cutoff for nsp12, 10 kDa for nsp8, and 3 kDa for nsp7.

### In vitro activity assays

The activity of the protein-RNA complex was assessed using in vitro extension assays. The polymerase was first assembled in a ratio of 5:5:1 for nsp12:nsp8:nsp7, 0.5 mg/mL nsp12 (equivalent to 5 µM), 0.5 mg/mL nsp8 (equivalent to 23 µM) and 0.25 mg/mL nsp7 (equivalent to 25 µM) before being incubated with pre-annealed RNA for 10 mins at 30°C in a 3 µL reaction containing master mix (MM) (15 mM MgCl2, 1.5 mM NTPs, 30 µM KCl, 3 U/µl RNasin, 0.3 mM DTT) and protein storage buffer (PSB) (12.5 mM HEPES, 75 mM NaCl, 1 mM MgCl_2_, 0.5 mM DTT, 40% glycerol) where RdRp:PSB:MM:RNA 1:3:1:1. For remdesivir and suramin, compounds were resuspended in diethylpyrocarbonate (DEPC)-treated water before being diluted to the final concentrations stated in the figures in PSB, replacing PSB during incubation with protein-RNA complexes. For denaturing gels, samples were denatured by incubation with loading dye (80% formamide, 10 mM EDTA) at 95°C for 3 minutes before being loaded onto a 6 M urea, 20% polyacrylamide denaturing gel and run at 450 V for 2.5 hours. For native gels, this step was replaced by adding a loading solution of 50% glycerol 1:1 with protein-RNA complexes and use of 4-15% Bio-Rad Mini-PROTEAN TGX gels which can be run in sodium dodecyl sulphate (SDS)-free tris-glycine running buffer. Gels were visualised using a Bio-Rad Chemidoc imaging system in the Cy3 and Cy5 emission ranges.

### FRET positioning software (FPS) modelling

RNA was modelled using 3D models made on the RNA Composer 2^17,18^ website and subsequently imported into Pymol and merged with the SARS-CoV-2 RdRp structure PDB 6YYT^3^. Fluorescence dyes are attached by a 6-carbon linker to the selected nucleotides shown in the schematics for each experiment and are attached via the C5’ of the indicated sugar residue. We used FRET Positioning Software (FPS)^19^ which models the accessible volume (AV) of each dye by considering the linker length, dye dimensions and steric hindrance to produce an average dye position and dye clouds which can be plotted in Pymol to represent the available positions for each dye. The distance between the average dye positions was measured using the ‘measure distance’ function in Pymol and used to estimate the distance between dye pairs. Distance was converted into an estimated FRET efficiency using an R0 for the Cy3/Cy5 pair of 5.4 nm, as previously characterised^20–22^.

### Single-molecule fluorescence spectroscopy

The EIFlex confocal microscope (Exciting Instruments) was used for single-molecule FRET experiments using alternating-laser excitation (ALEX) to excite donor and acceptor fluorophores sequentially^23^. RdRp at a final concentration of 3.8 uM was incubated with fluorescently labelled RNA at a final concentration of 13.5 nM for 30 mins at 30°C in PSB before dilution to a final RNA concentration of ∼50 pM for confocal imaging and analysis. We used a 2-laser line set up with 30-40 mW lasers at 520 nm and 638 nm. The 520 nm laser was used at 0.22 mW power with alternation speeds of OFF: 0 μS, ON: 45 μS, OFF: 55 μS, while the 638 nm laser power was 0.16 mW and alternating at OFF: 50 μS, ON: 45 μS, OFF: 5 μS. The Exciting Instruments software was used to measure and evaluate the signals obtained.

### Fluorescence correlation spectroscopy

The same microscope and experimental configuration as described above was used for fluorescence correlation spectroscopy (FCS) measurements. Excitation was at 520 nm in continuous-wave fashion at 0.1 mW laser power. Photon-by-photon arrival times in the donor and acceptor channels were correlated using pulsed interleaved excitation analysis (PAM) with MATLAB^24^. Data in the manuscript were derived from autocorrelation in the green detection channel, where the confocal volume parameters were determined using rhodamine 6G and it’s known diffusion coefficient 414 m^2^/s^25^. The correlations over lag time were fitted using PAM and plotted in MATLAB.

### Data analysis

The Exciting Instruments pipeline assigns fluorescent photons to donor or acceptor fluorophores based on photon arrival time and the two characteristic ratios, fluorophore stoichiometry, S and apparent FRET efficiency, E.

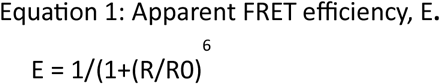

where E is the FRET efficiency, R is the inter-fluorophore distance and R0 is the Förster radius, a proportionality constant that depends on the interaction between the transition dipoles of donor and acceptor and represents the distance between the donor and the acceptor at which energy transfer is 50%.

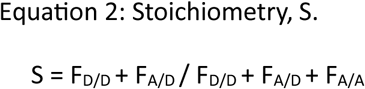

ALEX is used to alternately excite the donor (D) and acceptor (A) to determine the stoichiometry (S) of the molecules of interest taking into account signal (F) from the excited fluorophores. A two-dimensional histogram is plotted using S and E from fluorescent bursts above a certain threshold, and the one-dimensional E distribution of the populations within 0.4 < S < 0.8. We fitted these distributions with Gaussian functions to identify a mean E value for each distribution using the EI cloud software and associated Jupyter notebooks: FRETBursts (version 0.7.1)^26^. The notebooks performed an all photon burst search (APBS) and a dual channel burst search (DCBS)^27^ to collect bursts above a threshold, where selected photons passed through the detector at a rate 6x higher than background. The MultiFitter Gaussian model was used to fit 1 or more Gaussians, where the histogram and fit was displayed as a frequency (pdf=False) or probability density function (pdf=True).

## Results

### A minimal replication transcription complex is active in vitro

To establish a single-molecule SARS-CoV-2 RdRp replication assay we initially expressed and purified the minimal protein components needed for replication and transcription (the RTC): nsp12, nsp7 and nsp8, as well as a replication-deficient nsp12 mutant D760A/D761A (Fig 1A). The nsp12-His10-TEV-prot.A constructs were expressed in sf9 insect cells while the GST-3C-nsp7 and GST-3C-nsp8 plasmid constructs were expressed in E. coli under a lac promoter, before being purified using gravity-flow size-exclusion chromatography. Gel electrophoresis of the purified proteins showed bands at the expected sizes: 110 kDa for nsp12 wild type (WT) and nsp12 760A/D761A, 22 kDa for nsp8 and 9.4 kDa for nsp7 (Fig 1B).

**Figure 1:**
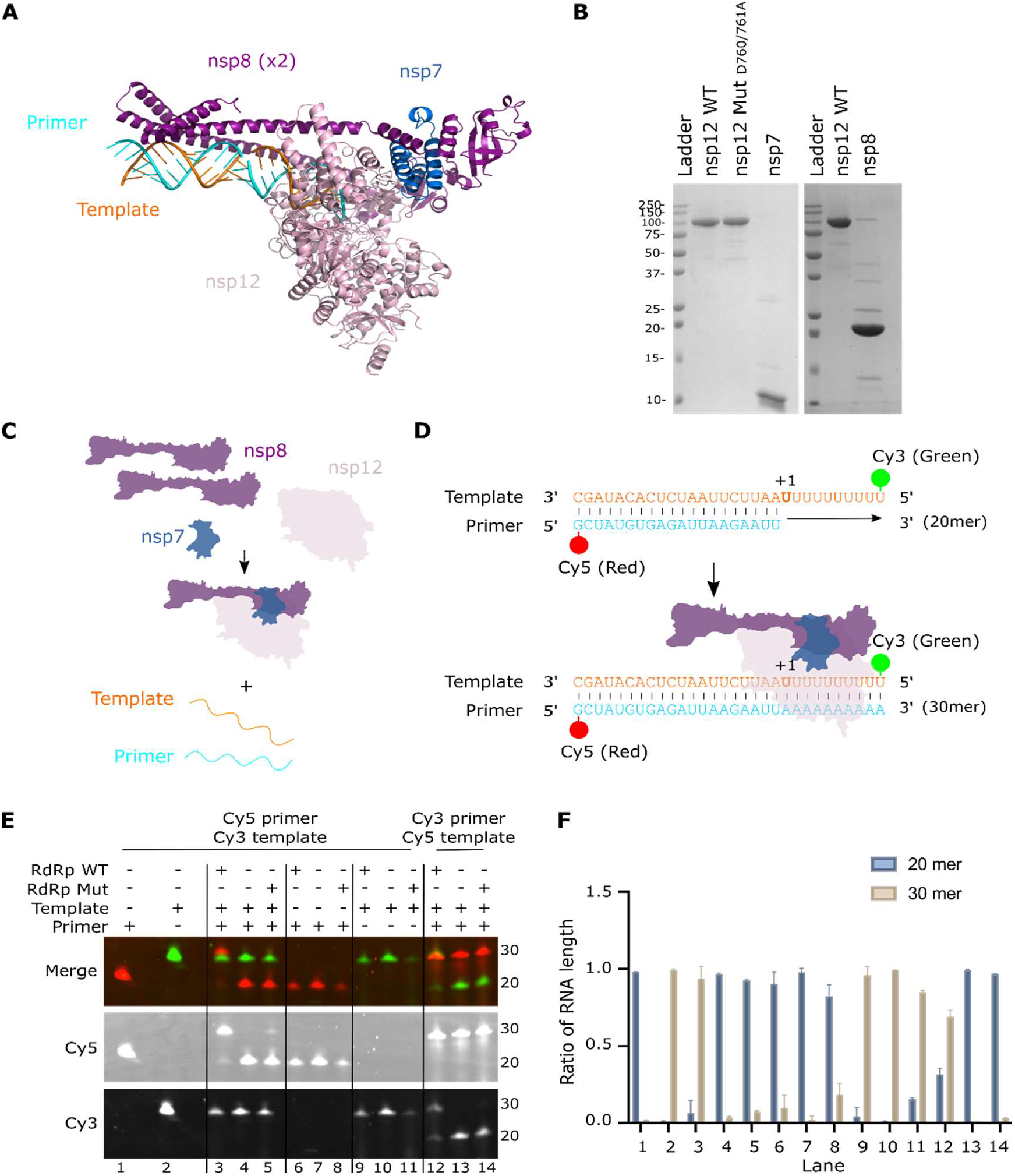
A minimal SARS-CoV-2 replication transcription complex (RTC) is active in vitro. A) Cryo-EM structure (PDB: 6YYT) of the SARS-C0V-2 RNA-dependent RNA polymerase bound to RNA. A single subunit of nsp12 (light pink) associates with two copies of nsp8 (purple) and one copy of nsp7 (dark blue), which in turn is bound to double-stranded RNA (orange template and cyan primer). B) SDS-PAGE gel showing the size and purity of the purified proteins; nsp12 wild type (WT), nsp12 active site mutant (D760A/D761A), nsp7 and nsp8. C) Schematic showing assembly of a minimal RTC by preincubation of the three nsps together to form a complex, before incubating with fluorescently labelled template and primer RNAs. D) Sequences of the annealed template (orange) and primer RNA (cyan), before and after RdRp-mediated extension. The template is labelled with Cy3 at position 1 and the primer is labelled with Cy5 at position 30. In the presence of RdRp and NTPs the primer sequence is extended to the same length as the template (30 mer). E) In vitro activity assay showing extension of the Cy5 labelled primer from a 20 mer to a 30 mer when RdRp WT, NTPs and both primer and template RNAs are present. F) Quantification of the relative products at the 20 mer position and 30 mer position in each lane of the gel, normalized to remove background fluorescence. Error bars represent standard deviation from two independent repeats.

To demonstrate activity of the purified RTC, we adapted a previous assay using ^32^P-radiolabelled RNA to verify polymerase activity^4,16^ for use with fluorescently labelled RNA substrates. We preincubated the three nsps together to form a complex, before incubating with fluorescently labelled template and primer RNAs (Fig 1C). Confirmation of activity is provided by extension of a short primer labelled with Cy5 from 20 to 30 nucleotides when RdRp is added (Fig 1D). Using the assay, we were able to show that a Cy5-labelled primer strand was only extended when the polymerase was complexed with both primer and template RNA strands (Fig 1E, Lane 3). We have quantified this as a ratio of RNA extension from a 20mer to 30mer using the intensity of the extended band over the total intensity of all bands, showing that the majority of RNA is extended in this assay if polymerase and both strands of RNA are present (Fig 1F). A Cy3-labelled primer was also extended in our assay, showing that in vitro activity is independent of the fluorescent dye (Fig 1E, Lane 12). As expected, the nsp12 active site mutant (D760A/D761A) demonstrated significantly lower activity than WT (Fig 1E, Lane 5). Overall, these results demonstrate that a purified minimal RTC was able to efficiently extend RNA labelled with fluorescent dyes.

### A single-molecule extension assay detects conformational changes in RNA during replication

To measure RdRp-mediated replication using smFRET, we used a 43-nucleotide ‘overhang’ RNA template that was designed to form a self-annealed hairpin labelled with a Cy3 dye at the 5’ end (Fig 2A). This template RNA was annealed to a shorter primer RNA, labelled with a Cy5 dye at position 12. Together, this template and primer RNA form a double-stranded construct that is predicted to bring the two dyes into close proximity, resulting in a high FRET signature. We hypothesised that upon RdRp and nucleotide addition the primer sequence would be extended, which in turn would open the self-annealed section of the template strand, resulting in the two dyes being separated and a decrease in the FRET signature (Fig 2A). Our template design also ensures that extension will proceed in a nucleotide-dependent manner, with partial extension expected from addition of ATP only, whilst full extension will only be achieved when both ATP and uridine triphosphate (UTP) are added. To estimate the efficiency of energy transfer between the two fluorophores in our expected constructs we used FRET Positioning Software (FPS) to calculate the accessible volumes that the Cy3 and Cy5 dyes could occupy when bound to the RNA in each configuration. We then measured the distance between the accessible volumes of the two dyes both for the RNA alone or RNA fully extended by the RdRp (Fig 2B). This measured distance allowed us to estimate the FRET efficiency for each dye pair, providing us an estimated FRET efficiency, E, of ∼0.99 for the RNA only and of ∼0.03 when the RNA is extended by polymerase. We therefore expect a switch from a high FRET efficiency to a low FRET efficiency once the RNA has been extended in vitro.

**Figure 2.**
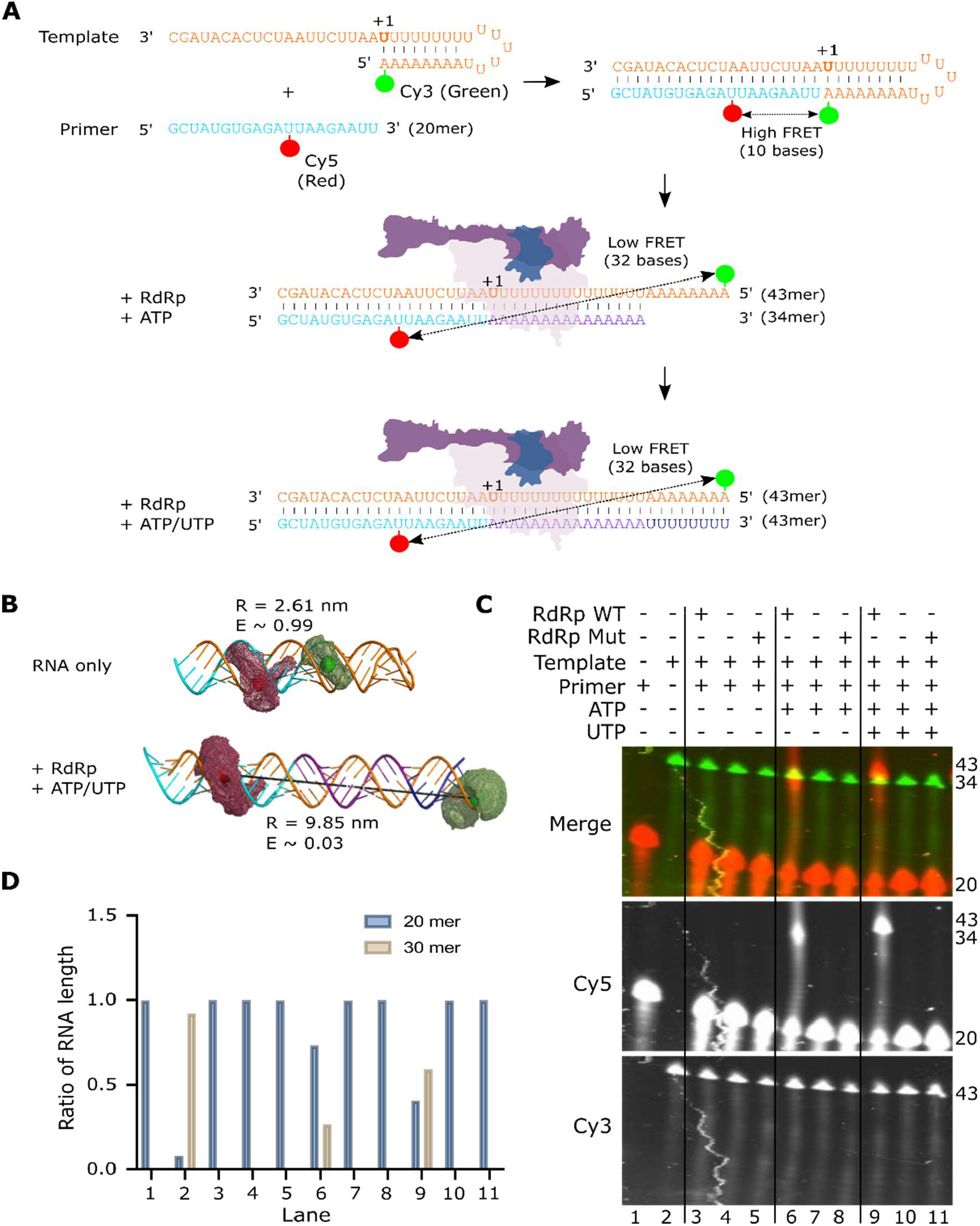
Design of a single-molecule extension assay to detect conformational changes in RNA during replication. A) Schematic showing the design of a single-molecule extension assay. A 43-nucleotide extension RNA template designed to form a self-annealed hairpin labelled with a Cy3 dye at the 5’ end was annealed to a short 20-nucleotide primer sequence, labelled with a Cy5 dye at position 12 (high FRET signature). Upon RdRp and nucleotide addition the primer sequence is extended, separating the two dyes (low FRET signature). B) Accessible volume modelling of the fluorescent dyes for non-extended and fully extended RNA. For non-extended RNA the dyes are 2.61 nm apart, giving an estimated FRET, E of 0.99, whilst the extended RNA dyes are an estimated 9.85 nm apart, giving an E ∼ 0.03. C) In vitro activity assay showing extension of the Cy5 labelled primer from a 20 mer to a 34 mer or 43 mer when RdRp and ATP or ATP/UTP were added. D) Quantification of the relative products at the 20 mer position and 34 or 43 mer position in each column, normalized to remove background fluorescence.

To confirm that our overhang RNA was active in vitro and that our new dye labelling positions didn’t reduce activity, we performed a fluorescence in vitro activity assay. We found that the primer in our replication complex was extended from a 20mer to a 34mer upon addition of RdRp WT and ATP alone, whilst addition of RdRp WT and both ATP and UTP resulted in full extension of the primer from a 20mer to a 43mer (Fig 2C). We quantified this as a ratio of RNA extension from 20mer to either a 34mer or 43mer using the intensity of each band over the total intensity (Fig 2D), which confirmed that the dsRNA construct can be extended in a nucleotide-dependent manner in the presence of WT polymerase.

Next, we used our overhang RNA construct to measure in vitro RdRp-extension using a diffusion-based smFRET assay (Fig 3 and Sup Fig 1A-E). We incubated the RNA with RdRp and nucleotides before diluting them to picomolar concentrations and measuring smFRET on replication complexes diffusing through the confocal volume. We found that annealed RNA alone produced a high FRET population centred at E = 0.90, confirming that the RNA forms the overhang RNA structure expected from our design (Fig 3A). As a control, we annealed a double-stranded RNA that mimics the expected product predicted from full extension of the primer and found that, as expected, this produced a significantly lower FRET population with mean E = 0.19 (Fig 3B). Addition of RdRp and ATP only resulted in two distinct FRET populations, a high FRET population representing non-extended overhang RNA (E = 0.90) and a low FRET population which represents partially extended RNA (E = 0.22) (Fig 3C). Addition of RdRp and both ATP and UTP results in full extension of the RNA primer, resulting in a major FRET population centred at E = 0.18, representing fully extended RNA, and a minor population of non-extended overhang RNA (E = 0.90) (Fig 3D). The assay using both ATP and UTP with the active site mutant nsp12 (D760A/D761A) showed no extension of the RNA, resulting in a major RNA population centred at E = 0.90 (Fig 3E). We also confirmed that we were able to get similar results with a different dye labelling position. We showed that moving the Cy5 dye from position 12 to position 19 on the primer abrogated extension activity (Sup Fig 2A-C), possibly because the dye prevented RdRp binding or was too close to the active site, however extension was observed when the position of the Cy3 dye on the template was changed from position 1 to position 3 (Sup Fig 2D-F).

**Figure 3.**
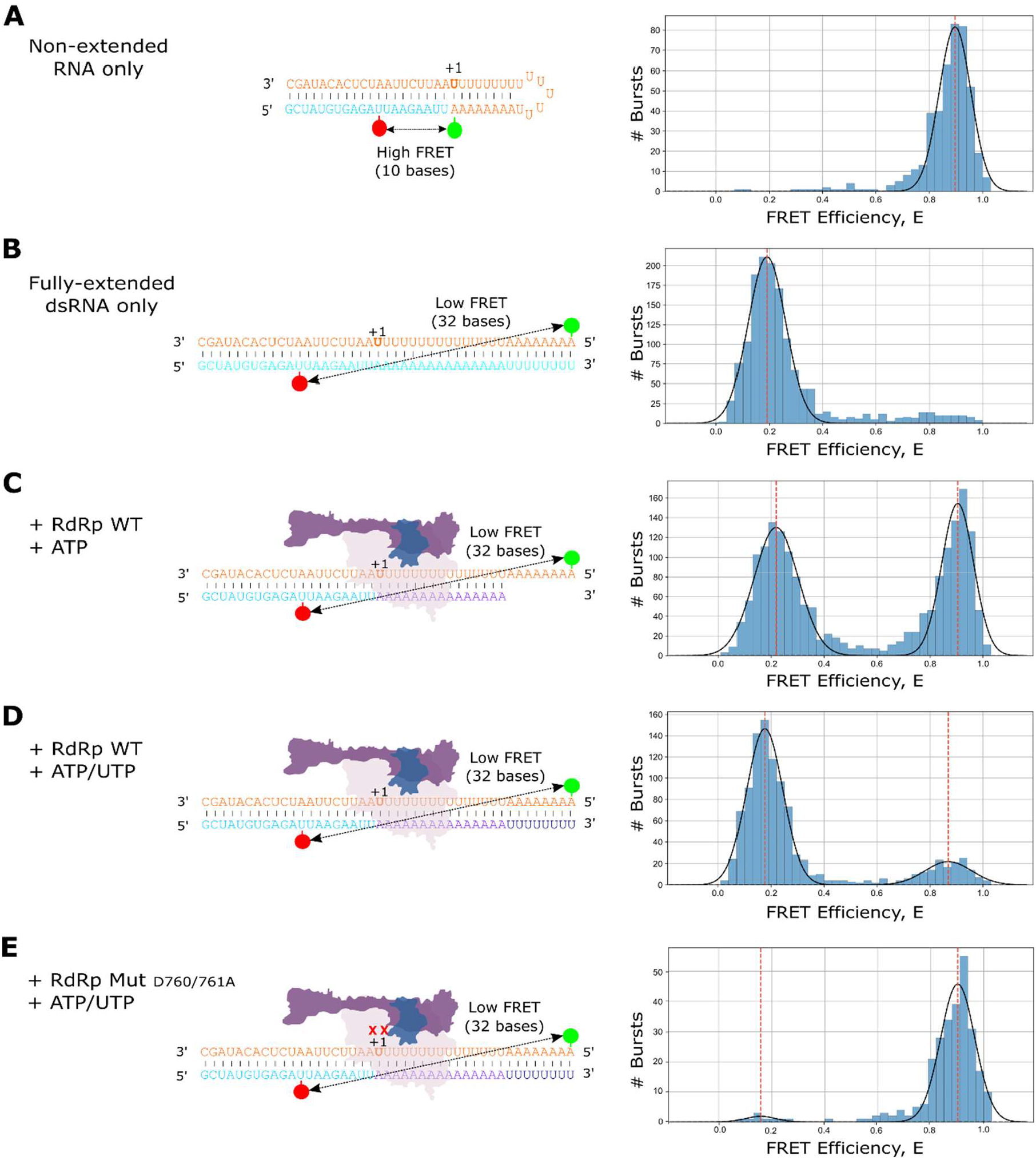
Single-molecule FRET can be used to measure RNA extension by the SARS-CoV-2 RdRp. A) FRET efficiency (E) when a Cy3-labelled 43mer template and Cy5-labelled 20mer primer were annealed without RdRp. B) A double-stranded RNA control that mimics the expected product predicted from full extension of the primer gives an expected low FRET population. C) Addition of RdRp and ATP to the pre-annealed RNA results in two distinct FRET populations (non-extended and partially extended RNA). D) Addition of RdRp and ATP/UTP extends the 20mer primer to a 43mer, resulting in almost complete RNA extension. E) Same as D), except using the RdRp mutant with nsp12 D760A/D761A (represented by XX in the figure).

Interestingly, we observed that addition of ATP only resulted in a lower conversion of overhang RNA to extended RNA compared to addition of both ATP and UTP (Sup Fig 1F). We hypothesised that addition of ATP only may lead to stalling of the RdRp on the partially extended replication complexes, therefore leaving fewer RdRp molecules available to replicate the RNA in the sample. In contrast, addition of both ATP and UTP should result in run-off of the RdRp from the end of the extended RNA, resulting in a readily available RdRp population that can further replicate any free RNAs in the sample. We confirmed our hypothesis by analysing the replication complexes on a native gel, which allowed us to see the complexes formed. The double-stranded RNA control that mimics the expected product predicted from full extension of the primer provided a marker for the expected size of the fully extended RNA complex (Sup Fig 3, lane 2), whilst addition of RdRp resulted in a larger complex that represents polymerase-bound RNA (Sup Fig 3, lane 3). As expected, addition of RdRp and ATP only led to increased stabilisation of the RdRp-RNA complex (Sup Fig 3, lane 4), whilst addition of RdRp with ATP and UTP resulted in a similar amount of complex to lane 3 (Sup Fig 3, lane 5), confirming that run-off of the RdRp occurs when the RNA is fully extended. Overall, our results indicate that the movements of RNA that occur during SARS-CoV-2 RdRp-mediated RNA extension can be efficiently captured by single-molecule FRET assays, and that these results are consistent with in vitro activity assay observations.

### Single-molecule FRET shows inhibition of RdRp by remdesivir

Having shown that our assay was able to capture RdRp-mediated RNA extension at the single-molecule level, we next sought to investigate the molecular mechanisms of replication inhibition. Remdesivir triphosphate is a licensed adenosine triphosphate analogue shown previously to inhibit SARS-CoV-2 RdRp. The proposed mechanism for remdesivir inhibition of RdRp is by delayed chain termination, whereby remdesivir triphosphate binds in place of ATP to hydrogen bond with UTP, and 3 more nucleotide triphosphates (NTPs) are added to the nascent strain before steric hinderance prevents further translocation and induces RdRp stalling (Fig 4A). We used a denaturing gel to show RdRp extension of a primer RNA in the presence of increasing concentrations of remdesivir with either high NTP (3mM final ATP/UTP) or low NTP (0.3mM final ATP/UTP) concentrations. Surprisingly, in high NTP conditions, we observed an increase in the fully extended 43mer product with increasing remdesivir concentration, suggesting that ATP and UTP are still being incorporated and that remdesivir itself may be mis-incorporated into the growing product (Fig 4B). We do observe some chain termination with 5 mM remdesivir, although a strong band representing the full-length product is also present. In low NTP conditions, significantly less full extension is observed and in the 5 mM remdesivir condition only partially extended primer products are produced (Fig 4B).

**Figure 4.**
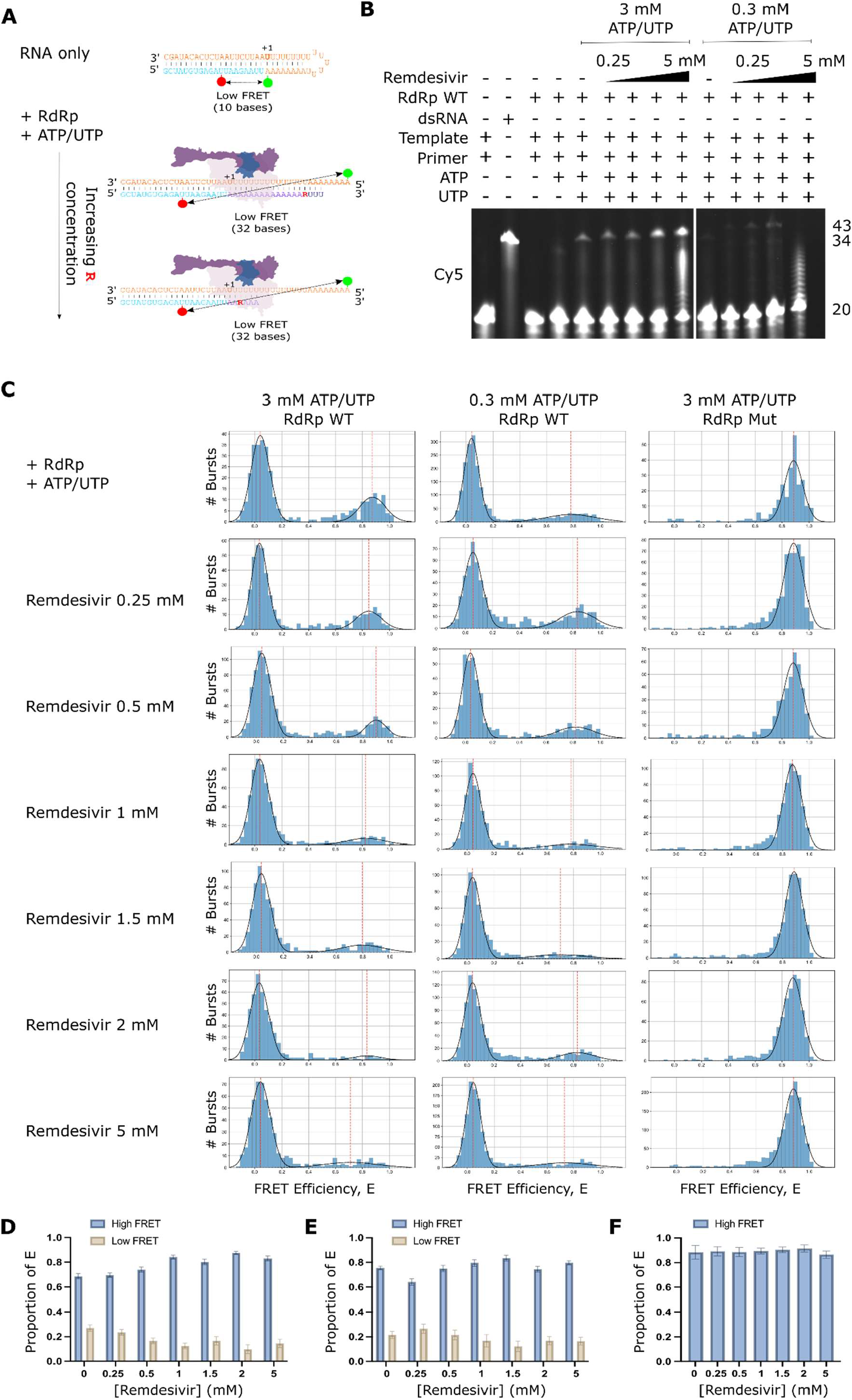
Inhibition of RNA extension with remdesivir. A) Schematic of the proposed delayed chain termination mechanism of action of the ATP analogue remdesivir (denoted by red ‘R’). B) Denaturing gel showing RdRp extension of a Cy5-labelled primer in the presence of ATP and UTP in increasing concentrations of remdesivir with either a high nucleotide triphosphate (NTP) (3 mM final) or low NTP (0.3 mM final) concentration. C) FRET efficiencies, E, for RNA conformations during extension by the RdRp in high NTP (3 mM final) concentration conditions or low NTP conditions (0.3 mM final), with either WT or mutant D670/671A RdRp. D-F) Quantification of the high and low FRET populations in C) as a proportion of the total FRET distributions by finding the probability density function. Error bars represent standard error of the gaussian fit.

Having established that remdesivir has an inhibitory effect on RdRp extension of our overhang RNA, we went on to image complexes using confocal smFRET. As expected from our denaturing gel results, we observed that in high NTP conditions, increasing concentrations of remdesivir resulted in an increase in the low FRET population representative of extended RNA, and a corresponding decrease in the high FRET population representative of non-extended overhang RNA (Fig 4C&D). Unexpectedly, however, we observed a similar pattern in low NTP concentration conditions (Fig 4C&E), compared to our gel results which suggested that we should observe a greater population of non-extended overhang compared to extended RNA. Exchange of the WT RdRp for the D760A/D761A mutant confirmed that extension was reliant on having an active RdRp present (Fig 4C&F).

### RdRp stalling plays a role in remdesivir inhibition

To further investigate our low NTP concentration smFRET results, we used a 4-15% gradient native gel to observe RdRp-RNA complexes with increasing concentrations of remdesivir, in both high (3 mM) and low (0.3 mM) NTP concentrations (Fig 5A&B). We incubated RdRp with RNA and 3 mM ATP/UTP and observed a strong band corresponding to the fully extended double-stranded RNA product on the gel resulting from run-off of the RdRp from the RNA, as well as a minor shifted band representing RNA that remained bound by the RdRp (Fig 5B, lane 1). Addition of remdesivir resulted in large complexes representing partially extended, RdRp-bound, RNA products (Fig 5B, lanes 2-5). Increasing concentrations of remdesivir resulted in an increasingly shifted band, suggesting that the RdRp was increasingly stalled on the increasingly extended RNA as remdesivir was incorporated. In low NTP conditions (0.3 mM ATP/UTP), no visible fully extended RNA product was observed (Fig 5B, lane 6), demonstrating that NTPs were limiting and full-length extension was inhibited. We expected that limiting NTP conditions would force remdesivir incorporation and chain termination, and consistent with this expectation, found that even low concentrations of remdesivir resulted in significant RdRp stalling on the RNA (Fig 5B, lanes 7-10). These results suggest that the extent of RdRp stalling on the viral RNA is heavily dependent on the concentration of natural NTPs.

**Figure 5.**
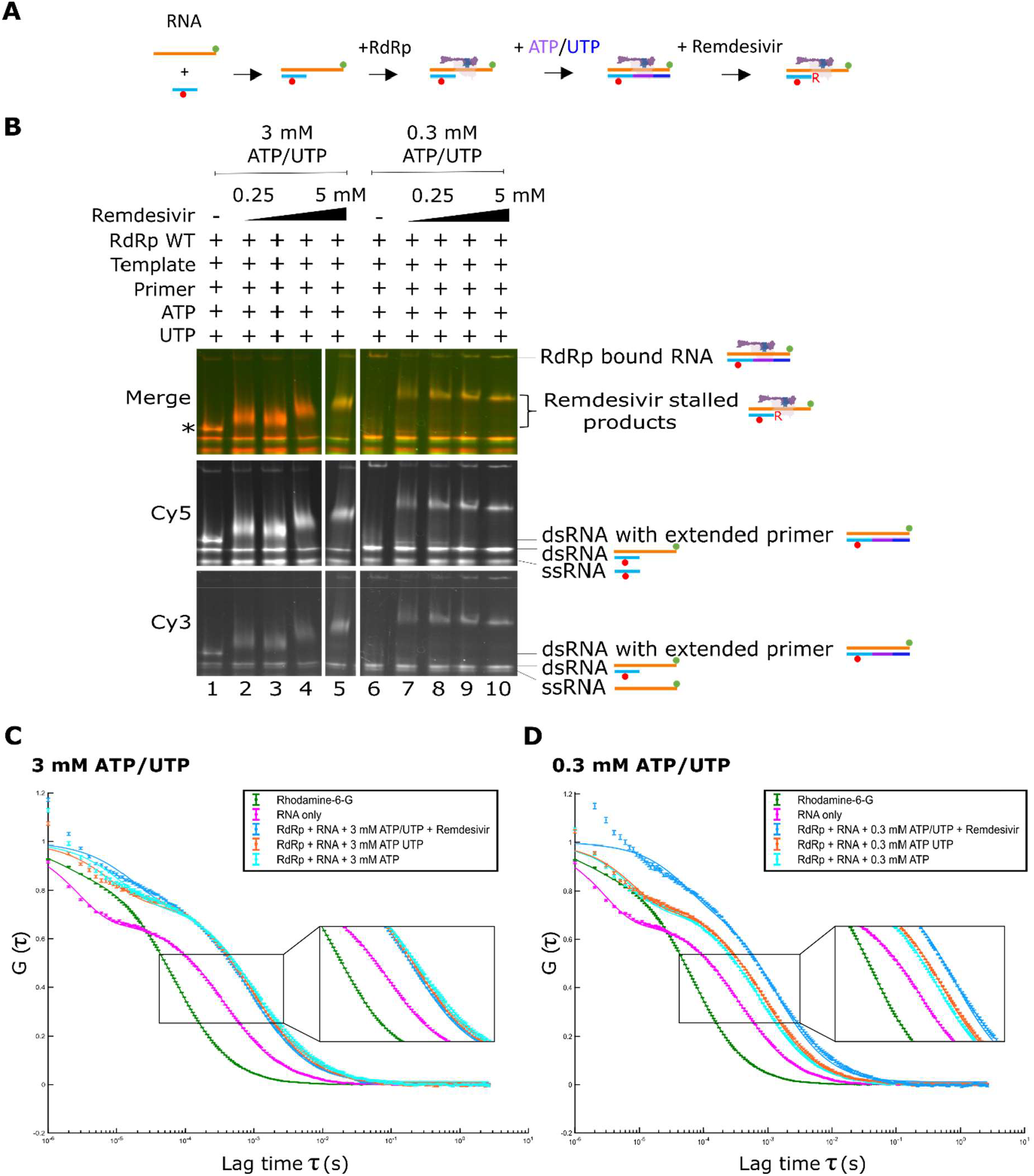
RdRp stalls on RNA in low NTP conditions. A) Schematic of RdRp binding to the fluorescently labelled RNA during NTP and remdesivir addition. B) Native gel showing remdesivir-enabled RdRp stalling on RNA in the presence of either 3 mM or 0.3 mM final NTP concentrations. * denotes the position of the extended double stranded RNA. C) Fluorescence correlation spectroscopy (FCS) comparing the diffusion of RNA only with RdRp-bound RNA in the presence of a 3 mM final concentration of nucleotides and remdesivir. Rhodamine 6G was used as a calibration reference. D) Fluorescence correlation spectroscopy (FCS) comparing the diffusion of RNA only with RdRp-bound RNA in the presence of a 0.3 mM final concentration of nucleotides and remdesivir.

To further investigate the role of RdRp stalling in SARS-CoV-2 replication inhibition by remdesivir we used fluorescence correlation spectroscopy (FCS), a method that measures the fluorescence intensity fluctuations of diffusing complexes, providing information on their diffusion rates using temporal autocorrelation. Rhodamine 6G was used as a standard to quantify the confocal volume parameters before RNA only or RNA-protein complexes were measured in either high (3 mM; Fig 5C) and low (0.3 mM; Fig 5D) NTP concentrations, and normalised correlations were plotted against lag time. At 3 mM ATP and UTP concentrations we observed a shift to longer lag times in all cases where RdRp was present compared to RNA only, as expected from the slower diffusion of larger complexes (Fig 5C). The addition of remdesivir, however, did not result in an additional increase in the diffusion time of the RNA in non-limiting natural NTP conditions (Fig 5C). In comparison, when remdesivir was added to the reaction with a final concentration of just 0.3 mM ATP and UTP, the diffusion time increased compared to similar conditions without remdesivir present (Fig 5D), supporting our other observations that there is increased stalling of the RdRp on the RNA when remdesivir is incorporated in low NTP conditions. We therefore conclude that in low NTP conditions remdesivir acts to inhibit the SARS-CoV-2 RdRp by delayed chain termination brought about by stalling the RdRp on the viral RNA, whilst in high NTP conditions the RNA can still be fully extended; the mechanism of action of remdesivir is therefore not only dependent on inhibitor concentration but also on the concentration of competing nucleotide triphosphates.

### Single-molecule FRET shows inhibition of RdRp by a non-nucleoside analogue inhibitor

To test inhibition of the SARS-CoV-2 RdRp by a non-nucleoside analogue inhibitor we assessed the impact of suramin addition to our assay. Suramin is a broad-spectrum inhibitor of viruses and parasites and has been shown to inhibit SARS-CoV-2 infection in cell culture by preventing cellular entry of the virus^28^ as well as inhibiting the viral polymerase by binding at two sites on the protein and blocking the binding of the RNA template strand and preventing entry of product RNA into the active site^13^ (Fig 6A). A denaturing gel where increasing concentrations of suramin were added to the reactions showed that extension of the overhang RNA was inhibited in a concentration-dependent manner (Fig 6B), which was supported by a native gel showing that fully extended RNA decreases in a similar way (Fig 6C). These results were supported by our single-molecule FRET experiments, in which increasing concentrations of suramin resulted in a decrease in the low FRET population representative of extended RNA (Fig 6D). Quantification of the ratios of low FRET and high FRET distributions supported this finding (Fig 6E). In summary, we have shown that single-molecule FRET can be a useful method to assess inhibition of the SARS-CoV-2 RdRp by a non-nucleoside inhibitor.

**Figure 6:**
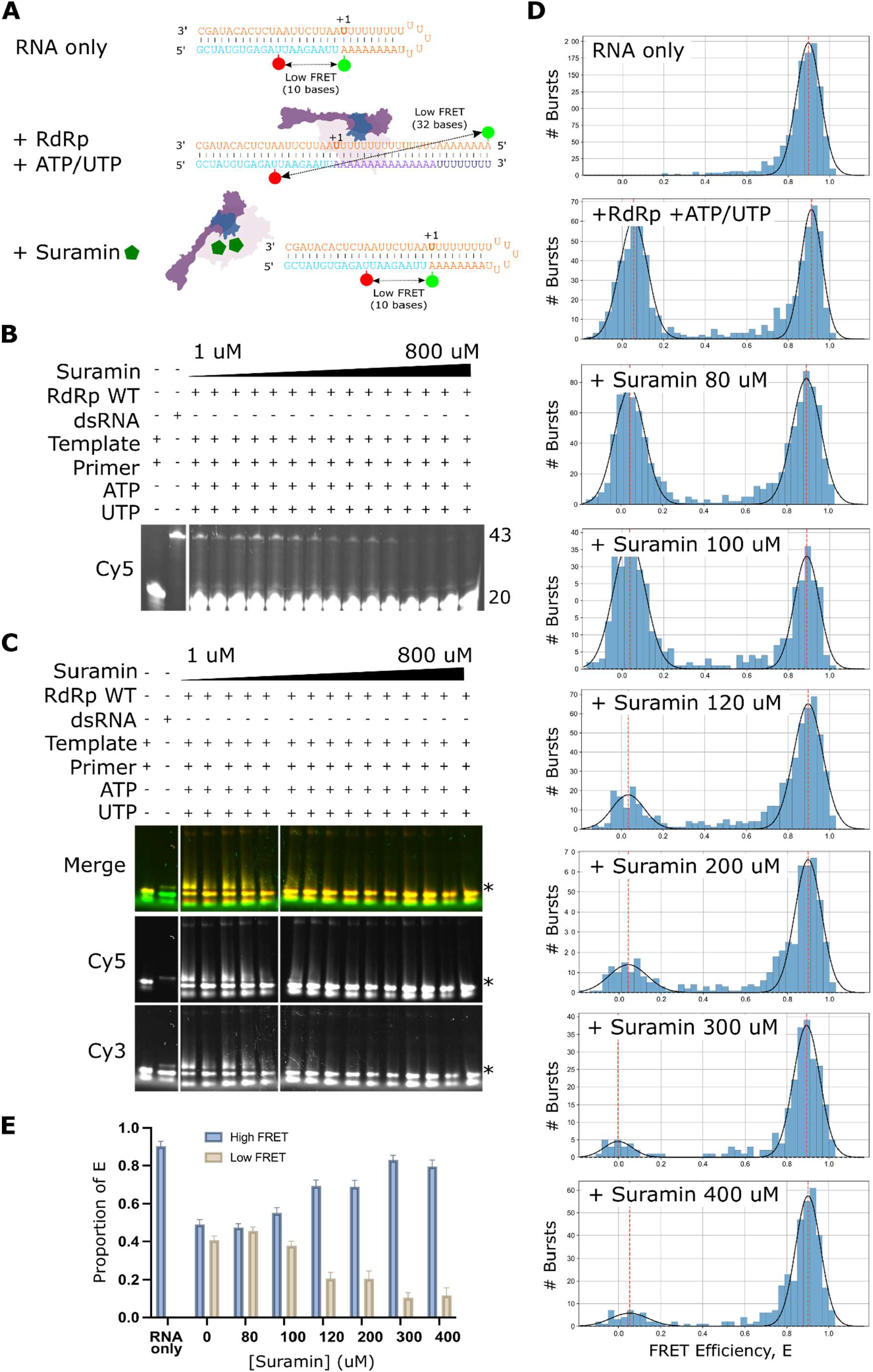
Single-molecule FRET analysis of RdRp inhibition by the non-nucleoside inhibitor suramin. A) Schematic of the proposed mechanism of action of suramin. The binding of two suramin molecules blocks the binding of the RdRp to the RNA template–primer duplex as well as the entry of nucleotide triphosphates into the catalytic site. B) Denaturing gel showing inhibition of RdRp extension of a Cy5-labelled primer in the presence of ATP and UTP in increasing concentrations of suramin. C) Native gel showing inhibition of fully extended primer product in increasing concentrations of suramin. * denotes the position of the extended double stranded RNA. D) FRET efficiencies, E, for RNA conformations during extension by the RdRp in increasing concentrations of suramin. E) Quantification of the high and low FRET populations in C) as a proportion of the total FRET distributions. Error bars represent standard deviation of the gaussian fit.

## Discussion

Despite the devastating impact of the COVID-19 pandemic, many of the molecular details of coronavirus replication remain unknown. Here, we have expressed and purified the minimal protein components of the SARS-CoV-2 replication complex and shown that these are capable of actively extending RNA labelled with fluorophores. Although this complex has previously been shown to extend radiolabelled RNA^4^, we have developed fluorescence-based extension assays to test RdRp activity without the need for radioactive components. This novel approach has allowed us to study SARS-CoV-2 minimal replication complexes using fluorescence-based in vitro activity assays and single-molecule imaging. To measure RdRp-mediated replication using smFRET, we used a unique ‘overhang’ RNA template that was designed to form a self-annealed hairpin that upon nucleotide addition to the primer strand would open and cause a decrease in the expected FRET signature. We were also able to control nucleotide addition through the template design, ensuring that either partial or full RNA extension took place. Our assays allowed us to observe SARS-CoV-2 RdRp-mediated extension of RNA at the single-molecule level, and to assess the impact of both non-nucleoside and nucleoside analogue inhibitors on extension.

Remdesivir is a direct-acting antiviral agent that is used to treat patients with severe COVID-19 disease by targeting the viral RdRp of SARS–CoV-2. Despite having broad-spectrum antiviral activity against a wide variety of viruses the mechanisms of action of remdesivir aren’t fully understood. The triphosphate form of remdesivir competes with its natural counterpart adenosine triphosphate (ATP) for incorporation into viral RNA, thought to lead to delayed chain termination^8,10,11,29^. Our gels and single-molecule FRET experiments showed, however, an increase in the fully extended RNA product with increasing remdesivir concentration when the natural ATP and UTP were at a high concentration of 3 mM. This is in agreement with a previous observation that RNA synthesis arrest can be overcome with higher concentrations of natural nucleotide pools^11^. Interestingly, intracellular NTP concentrations are in the high μM and low mM range^30^, which suggests that read-through reactions likely occur under biologically relevant conditions. If read-through reactions occur in a cellular context it is likely that remdesivir itself may be mis-incorporated into the growing product. It has therefore been suggested that the primer strand, and by extension the negative-sense copy of the viral genome, could contain several remdesivir residues^12^. Inefficient incorporation of UTP opposite remdesivir molecules in the genome may therefore provide a second explanation for how remdesivir inhibits SARS-CoV-2 replication^12^.

In contrast to remdesivir inhibition in high NTP conditions, our gels showed that significantly less full extension was observed when natural ATP and UTP were at a lower concentration of 0.3 mM. This result wasn’t reflected in our single-molecule FRET experiments, however, which showed a similar pattern to that observed in high NTP concentration conditions, where a greater population of fully extended RNA was observed than non-extended overhang RNA. This was explained by using native gel electrophoresis and FCS to show that in limiting natural NTP conditions increasing concentrations of remdesivir resulted in increasingly more protein stalling on the RNA. Stalled RdRp would be expected to sit on the RNA template and hold it in an extended conformation, thus resulting in a low FRET signature. Our results are consistent with other findings, for instance studies that provided a structural analysis of RdRp stalling by remdesivir^31,32^ and showed that RdRp translocation is prevented by the sterically impaired passage of the cyano group in remdesivir past the serine-861 side chain in the nsp12 subunit of RdRp. A second study that used a single-molecule magnetic-tweezers platform also showed that the barrier induced by the clash of remdesivir with serine-861 was sufficiently strong to elicit polymerase stalling and backtracking^33^. Taken together, this suggests that the predominant mechanism of remdesivir action against the SARS-CoV-2 RdRp may be dependent on the surrounding concentration of competing nucleotide triphosphates.

Our fluorescence-based single-molecule FRET assays also proved to be a useful technique to assess inhibition of the SARS-CoV-2 RdRp by suramin, a non-nucleoside inhibitor. Suramin was first described over one hundred years ago, and has been shown to be effective at inhibiting the replication of a wide range of viruses, including enteroviruses, Zika virus, Chikungunya, Ebola and SARS-CoV-2^34^. A cryo-EM structure of suramin bound to the SARS-CoV-2 RdRp revealed that suramin bound to the RdRp active site, blocking the binding of both RNA template and primer strands. This mechanism of action was supported by our single-molecule FRET results, which showed that increasing concentrations of suramin resulted in a decrease in extended RNA.

The minimal RTC used in our experiments does not incorporate other important viral nsps involved in replication, such as the multifunctional nsp14 enzyme which has 3’-5’ exoribonuclease and methyltransferase activity^35^, or the helicase nsp13. Nevertheless, our findings are consistent with several other studies of SARS-CoV-2 viral replication and our simple system has the potential for additional nsps to be added to allow us to mechanistically study their function at the single-molecule level.

In summary, our results advance our understanding of the interactions of the SARS-CoV-2 RdRp with RNA and a variety of viral inhibitors. As the RdRp complexes of SARS-CoV, MERS-CoV, and SARS–CoV-2 have all been suggested to be inhibited by remdesivir via a delayed chain termination mechanism it is likely that all three of these dangerous viruses exhibit common mechanisms of inhibition; our results therefore provide the potential to study alternative mechanisms of action for other viral RdRps and inhibitors. Our unique single-molecule hairpin RNA design therefore not only offers a useful way to rapidly screen multiple compounds for potential antiviral activity, but also offers the ability to disentangle the exact mechanisms of inhibition of inhibitor compounds, all of which is crucial to allow us to screen for and design new generations of antivirals.

## Data availability

The data underlying this article are available in the article and in its online supplementary material.

