## Supplementary material for "Mechanistic insights into the activity of SARS-CoV-2 RNA polymerase inhibitors using single-molecule FRET"

| **RNA** | **Sequence** |
| --- | --- |
| Template | 5' - /Cy3/UUUUUUUUUUAAUUCUUAAUCUCACAUAGC - 3' |
| Primer | 5' - /Cy5/GCUAUGUGAGAUUAAGAAUU - 3' |
| Template swapped | 5' - /ATTO647N/UUUUUUUUUUAAUUCUUAAUCUCACAUAGC - 3' |
| Primer swapped | 5' - /Cy3/GCUAUGUGAGAUUAAGAAUU - 3' |
| Extension Template P1 | 5' - /Cy3/AAAAAAAAUUUUUUUUUUUUUUUAAUUCUUAAUCUCACAUAGC - 3' |
| Extension Primer P12 long | 5' - GCUAUGUGAGAU/Cy5/UAAGAAUUAAAAAAAAAAAAAAAUUUUUUUU - 3' |
| Extension Primer P12 short | 5' - GCUAUGUGAGAU/Cy5/UAAGAAUU - 3' |
| Unlabelled Template | 5' - UUUUUUUUUUAAUUCUUAAUCUCACAUAGC - 3' |
| Unlabelled Primer | 5' - GCUAUGUGAGAUUAAGAAUU - 3' |
| Unlabelled extension Template | 5' - AAAAAAAAUUUUUUUUUUUUUUUAAUUCUUAAUCUCACAUAGC - 3' |
| Template Cy3 P3 | 5' - CCA/Cy3/AAAAAAAUUUUUGGUUUAAUUCUUAAUCUCACAUAGC - 3' |
| Primer Cy5 P19 | 5' - GCUAUGUGAGAUUAAGAAU/Cy5/U - 3' |

**Supplementary Table 1.** Sequences and fluorescence dye positions of the RNAs used in the study. /Cy3/: position of the Cy3 label, /Cy5/: position of the Cy5 label, /ATTO647/: position of the ATTO647 label.

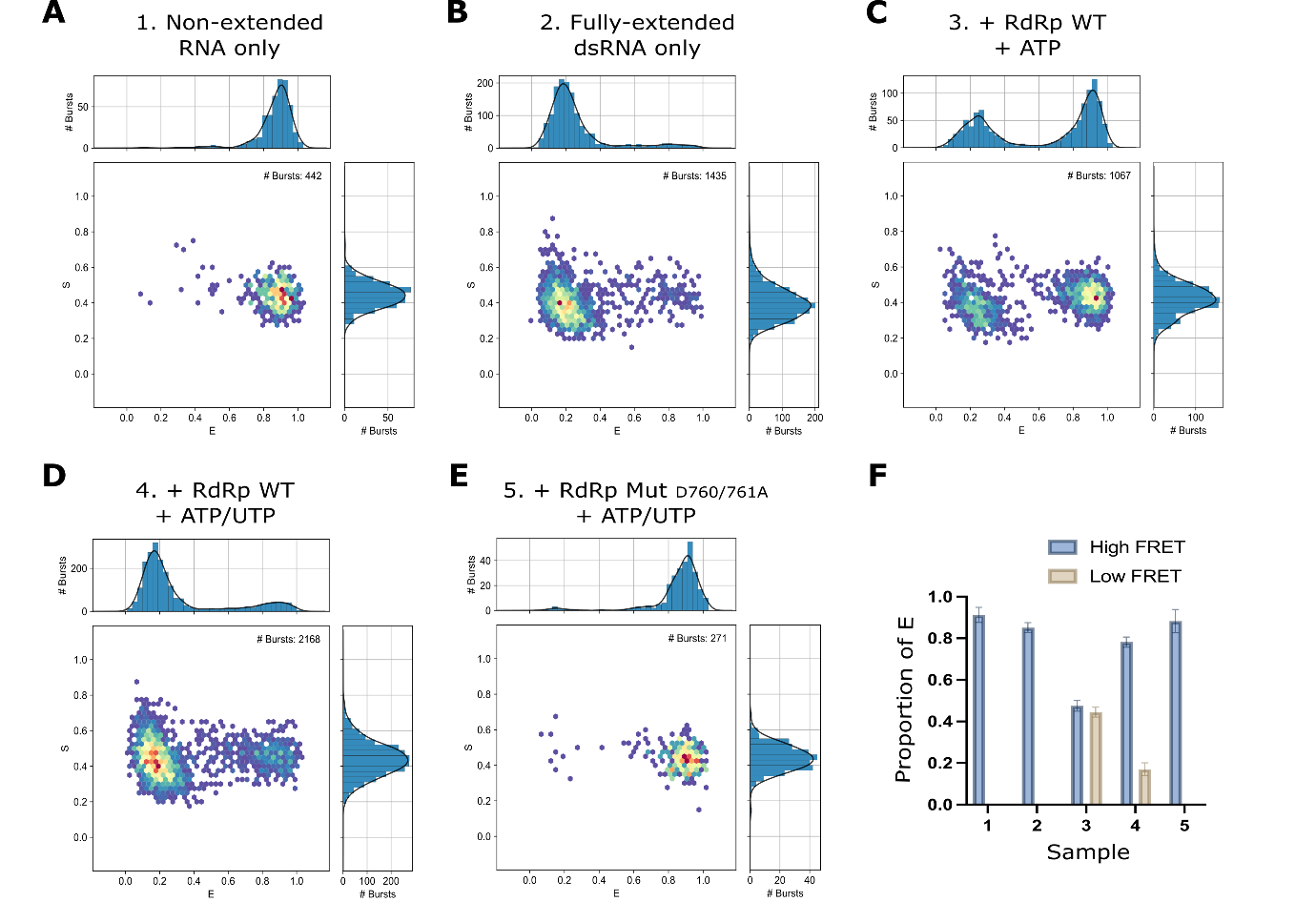

**Supplementary Figure 1. Single-molecule FRET can be used to measure RNA extension by the SARS-CoV-2 RdRp.** A-E) Single-molecule E-S histograms from the data presented in Fig 3. E represents apparent FRET efficiency and S represents stoichiometry. F) Quantification of the high and low FRET populations in A-E as a proportion of the total FRET distributions.

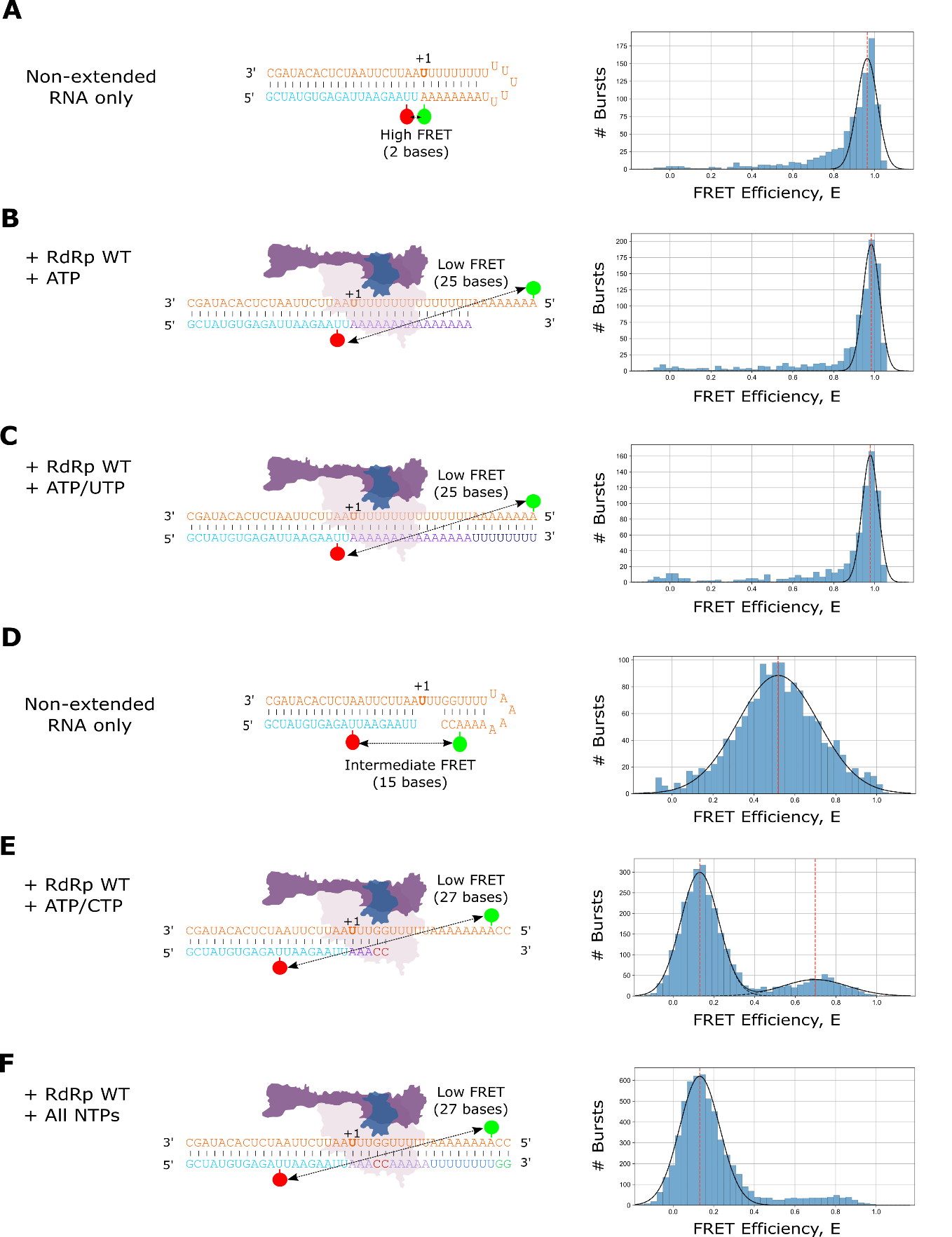

**Supplementary Figure 2. Investigation of alternative labelling positions for the single-molecule FRET extension assay.** A) FRET efficiency (E) for RNA only when the Cy5 dye on the 20mer primer was moved to position 19. B) Addition of RdRp and ATP to the pre-annealed RNA resulted in no extension of the primer, suggesting that the dye at position 19 is inhibitory. C) Addition of RdRp and ATP/UTP also resulted in no extension of the primer. D) FRET efficiency (E) for RNA only when the Cy3 dye on a shortened template was moved to position 3. E) Addition of RdRp and ATP/CTP to the pre-annealed RNA resulted in a shift to low FRET, suggesting opening of the template RNA. F) Addition of RdRp and all NTPS to the pre-annealed RNA provided similar results to E).

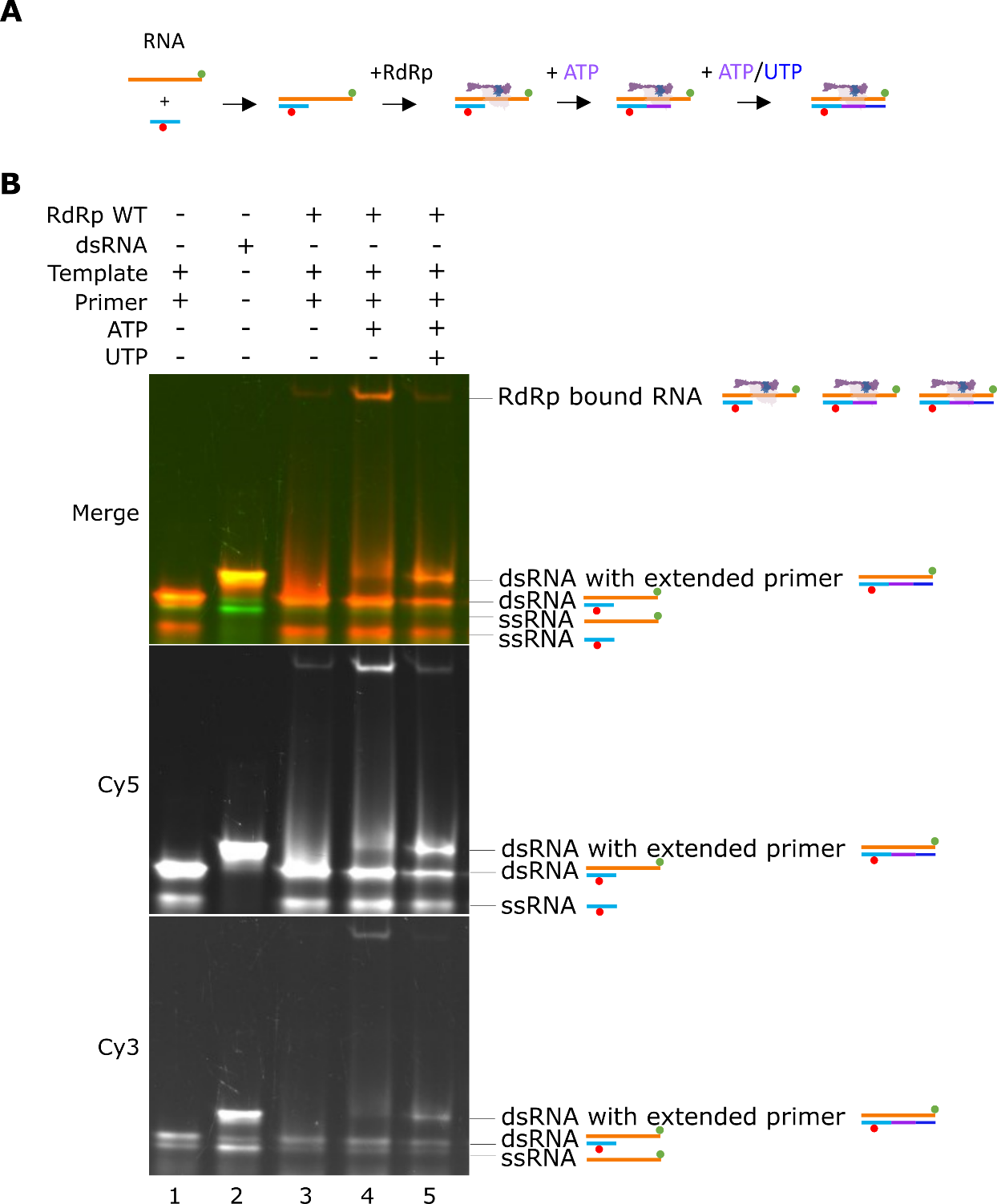

**Supplementary Figure 3. Native gel electrophoresis shows RdRp-RNA complexes during replication.** A) Schematic of RdRp binding to the fluorescently labelled RNA during sequential NTP addition B) Native gel showing binding of RdRp to the extension RNA. Labelled primer and template RNA run as two distinct single-stranded RNA bands as well as a double-stranded RNA (lane 1). A double-stranded RNA control that mimics the expected product predicted from full extension of the primer gives the position of the fully extended RNA (lane 2). Addition of RdRp alone does not result in primer extension but does show an RdRp-bound band higher in the gel (lane 3). Addition of RdRp and ATP results in enhanced stalling of RdRp on the partially extended primer substrate (lane 4), whilst addition of ATP and UTP results in an increase in run-off of the RdRp and an increase in the double-stranded RNA product (lane 5).
